# An ancient endogenous DNA virus in the human genome

**DOI:** 10.1101/2022.02.01.478760

**Authors:** Jose Gabriel Nino Barreat, Aris Katzourakis

**Author notes:** Address correspondence to Aris Katzourakis.

## Abstract

The genomes of eukaryotes preserve a striking diversity of ancient viruses in the form of endogenous viral elements (EVEs). Study of this genomic fossil record provides insights into the diversity, origin and evolution of viruses across geological timescales. In particular, *Mavericks* have emerged as one of the oldest groups of viruses infecting vertebrates (≥419 My). They have been found in the genomes of fish, amphibians and non-avian reptiles but had been overlooked in mammals. Thus, their evolutionary history and the causes of their demise in mammals remain puzzling questions. Here, we conduct a detailed evolutionary study of two *Maverick*-like integrations found on human chromosomes 7 and 8. We performed a comparative analysis of the integrations and determined their orthology across placental mammals (Eutheria) via the syntenic arrangement of neighbouring genes. The integrations were absent at the orthologous sites in the genomes of marsupials and monotremes. These observations allowed us to reconstruct a time-calibrated phylogeny and infer the age of their most recent common ancestor at 268.61 (199.70–344.54) My. In addition, we estimate the age of the individual integrations at ~105 My which represent the oldest non-retroviral EVEs found in the human genome. Our findings suggest that active *Mavericks* existed in the ancestors of modern mammals ~172 My ago (Jurassic Period) and potentially to the end of the Early Cretaceous. We hypothesise *Mavericks* could have gone extinct in mammals from the evolution of an antiviral defence system or from reduced opportunities for transmission in terrestrial hosts.

**Importance:** The genomes of vertebrates preserve an enormous diversity of endogenous viral elements (remnants of ancient viruses that accumulate in host genomes over evolutionary time). Although retroviruses account for the vast majority of these elements, diverse DNA viruses have also been found and novel lineages are being described. Here we analyse two elements found in the human genome belonging to an ancient group of DNA viruses called *Mavericks*. We study their evolutionary history, finding that the elements are shared between humans and many different species of placental mammals. These observations suggest the elements inserted at least ~105 Mya in the most recent common ancestor of placentals. We further estimate the age of the viral ancestor around 268 My. Our results provide evidence for some of the oldest viral integrations in the human genome and insights into the ancient interactions of viruses with the ancestors of modern-day mammals.

## Introduction

The genomes of vertebrates are teeming with viruses. Some of these elements are still active (they can replicate and produce viral progeny), while most are degraded genomic fossils. The most prominent are the retroviruses (RNA-retrotranscribing viruses) which can actively integrate and account for a significant proportion of host genomes (1). Remarkably, some animal DNA viruses also share this property of actively integrating into host genomes, including *Mavericks* (2, 3), *Teratorns* (4–6), herpesviruses (7–9) and adeno-associated virus (AAV) (10, 11).

*Mavericks* are a group of eukaryotic mobile genetic elements that encode a family B polymerase, a retroviral-like integrase and homologues of capsid morphogenesis genes found in dsDNA viruses (2, 3, 12, 13). Evolutionary analyses and comparisons of protein structure by homology modelling indicate that *Mavericks* are firmly nested within the viral kingdom *Bamfordvirae* (12–15). They are considered to be a lineage of endogenising DNA viruses that can undergo exogenous transmission (16, 17), although there is still no direct experimental evidence for their assembly into viral particles.

In vertebrates, *Mavericks* have been described in the genomes of teleost fish, coelacanths, amphibians, birds and non-avian reptiles. Most of the potentially active elements occur in teleost fish (97%) while only 4 intact copies (3%) have been found in tetrapod genomes (16). All the elements found in birds seem to be degraded and they had not been identified in mammals (2, 3, 16). The reason for the deterioration and eventual loss of the elements from birds and mammals remains unclear.

Here we report the discovery of two novel placental-wide *Maverick* orthologues also found in humans which allow us to conduct a detailed evolutionary analysis of the elements across mammals (human sequences on chromosome 7 have been identified in previous work (17), but their orthology and evolution have not been investigated). By comparing these loci and placing them in a phylogenetic context, we gain new insights into the evolutionary history of *Mavericks* in mammals and the forces that have shaped their evolution. Furthermore, we discuss several hypotheses which could explain the demise of the elements in bird and mammalian genomes. The findings presented here illuminate our understanding about the evolutionary history of *Mavericks*, the endogenous viral elements in the human genome and the ancient interactions between viruses and our direct ancestors.

## Methods

We used the protein-primed polymerase B (PolB) of the common box-turtle (*Terrapene carolina*) *Maverick* (16, 17) as a probe to screen the human genome with tBLASTn (18) (assembly GRCh38.p13, BLOSUM 45 substitution matrix, word size = 2). We obtained two hits with e-values < 1e-7: one on chromosome 7 (chr7:37601124–37601615, query cover = 15%, e-value = 8e-8, percent identity = 31.14%) and another on chromosome 8 (chr8:12858023–12858613, query cover = 25%, e-value = 5e-8, percent identity = 29.95%). In order to assess orthology of the integrations in the genomes of other mammals, we identified the three most proximal protein-coding genes upstream and downstream of the integrations using the Ensembl genome browser (19).

The protein-coding genes used as genetic landmarks for the chromosome 7 integration were: *NME8*, *EPDR1*, *GPR141*, *ELMO1*, *AOAH* and *ANLN*, and for chromosome 8: *C8orf48*, *DLC1*, *TRMT9B*, *LONRF1*, *FAM86B2* and *DEFB130A* (NCBI protein database (20)). We used these sequences as queries in tBLASTn (set to the default) to screen the NCBI RefSeq genome database using the following taxon labels: ‘Monotremata (taxid: 9255)’, ‘Metatheria (taxid: 9263)’, ‘Afrotheria (taxid: 311790)’, ‘Xenarthra (taxid: 9348)’, ‘Laurasiatheria (taxid: 314145)’ and ‘Euarchontoglires (taxid: 314146)’. To verify the presence or absence of the integration, we screened the regions flanked by the genetic landmarks with the full set of proteins coded by the 4 tetrapod *Mavericks* reported previously (16) again with tBLASTn (e-value < 1e-5).

As an independent test for the presence of the integrations, we used the whole-genome alignment of 120 mammals published by Hecker and Hiller (21). We extracted the regions corresponding to the human *polB* integrations and extended them assuming the ancestral gene coded for 1053 amino acids as in the box-turtle *Maverick* (chr7:37600749–37603904, chr8:12856919–12860047). The regions were extracted from the chr7 and chr8 whole-genome alignments in MAF format (85.16 GB and 81.07 GB, respectively) with custom *.bed files and using the mafsInRegion utility (22). Subalignments were transformed into fasta format with the MAF to FASTA programme (version 1.0.1) in Galaxy (23). The taxa present in these subalignments was then compared to those obtained with the BLAST-hit method.

We merged the chromosome 7 and 8 subalignments corresponding to the *polB* gene using the –merge function in MAFFT. The alignment was trimmed in trimAl version 1.4.rev22 (24) with the –automated1 option (which selects the optimal trimming method for the alignment). We then selected the best nucleotide substitution model in ModelTest-NG version 0.1.7 (25) (TVM+Γ4 under the AIC, BIC and cAIC criteria). Having determined the orthology of the integrations, we conducted a phylogenetic analysis under a relaxed-molecular clock using divergence-time calibrations in BEAST 2 version 2.6.6 (26). For this, we used 13 calibration points obtained from TimeTree (27) (Table A1) using log-normal distributions, and then ran a Bayesian MCMC chain for 100,000,000 generations, sampling every 5,000^th^ generation. We confirmed that the analysis had converged by inspecting the mixing and stationarity of posterior samples and ensuring that the Estimated Sample Sizes (ESSs) were greater than 200 for all parameters (burn-in of 25%).

Finally, we tested the possibility of selective constraints acting on the integrations. To this end, we calculated the pairwise genetic distances (measured as the observed proportion of nucleotide differences) between the taxa in the chromosome 7 and chromosome 8 subalignments separately, and compared these to 100 surrounding non-coding genomic regions of the same size in each case. We built two empirical distributions from these, and then tested the hypothesis that the distribution of *Maverick* genetic distances was distinct from the non-coding distribution. This was done with the a one-tailed, non-parametric Kolmogorov-Smirnov test. An R script was developed for this purpose using functionality from the Ape package (28). We further checked if the human integrations fell within piRNA clusters reported for chromosomes 7 and 8 (29), but this resulted in no matches (Supplementary excel file).

## Results

The *Maverick*-like integrations on chromosomes 8 and 7 are orthologous across the clade of placental mammals (Eutheria). The orthology of the regions was validated by the syntenic arrangement of the most proximal protein-coding genes, presence of the taxa in the whole-genome alignment of 120 mammals, and the relative arrangement of BLAST hits to different *Maverick* proteins which were consistent with the genetic organisation of vertebrate *Mavericks*. In addition, the taxonomic distributions resulting from the BLAST-hit and whole-genome alignment methods were consistent with each other (Supplementary excel file and multiple sequence alignments). However, some taxa were present in the whole-genome alignment corresponding to the *polB* marker which were not detected with the BLAST method, probably because of high divergence of the sequences (producing higher e-values than the cut-off). We could not detect orthologous integrations in the genomes of monotremes (*Ornithorhynchus anatinus*) or marsupials neither through BLAST nor in the whole-genome alignments.

Orthologues to the human chromosome 7 element were found in the genomes of primates, rodents, afrotherians (elephants, manatees and aardvark) and xenarthrans (sloths and armadillos) (Figure 1). In addition, the *polB* marker was present in the whole genome-alignment for scandentians (tree shrews), dermopterans (flying lemurs), lagomorphs (hares) and two laurasiatherians: the star-nosed mole (*Condylura cristata*) and the European hedgehog (*Erinaceus europaeus*). This finding is consistent with the presence of hits to the minor capsid protein (but not the polymerase) in other laurasiatherians, namely carnivores and cetaceans (Supplementary excel file). Remarkably, we detected an additional hit to the integrase in the genome of the brown bear (*Ursus arctos*): as expected, it occurs in the same reading frame as the minor capsid and does not show similarity to the integrases of retroelements in Censor (30). Orthologues to the human chromosome 8 integration were found in the genomes of primates and xenarthrans (Linnaeus’ two-toed sloth, *Choloepus didactylus*). The *polB* marker again had a broader taxonomic distribution in the whole-genome alignment and also included: scandentians, dermopterans, lagomorphs and chiropterans (bats) (Figure 2). In this case, we could not detect orthologues neither in rodents, afrotherians nor in other laurasiatherians.

**Figure 1.**
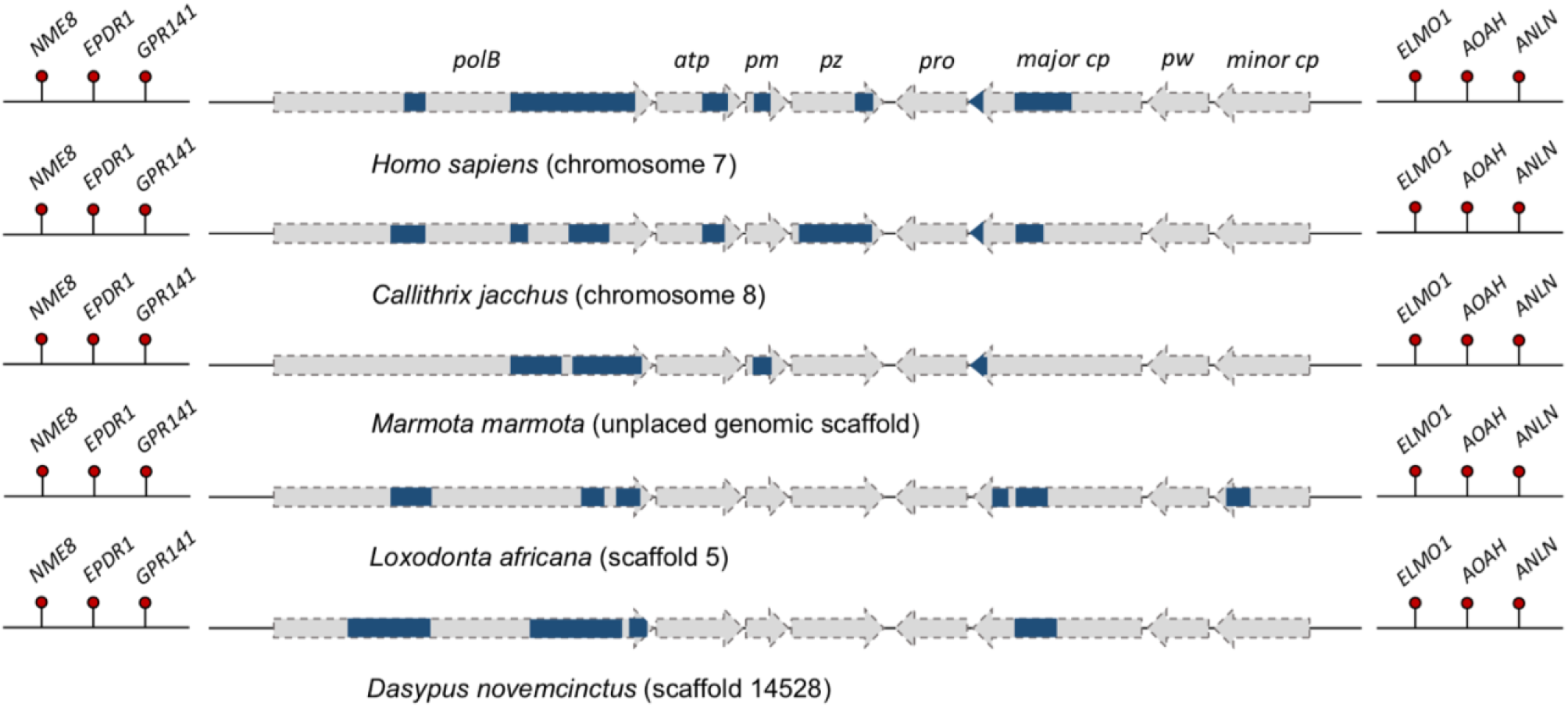
Comparison of the regions orthologous to the chromosome 7 *Maverick* integration in humans (*Homo sapiens*). tBLASTn hits (e-value < 0.05) to the proteins encoded by the common box-turtle (*Terrapene carolina*) *Maverick* are shown in blue. The arrow outlines represent the open reading frames of the box-turtle’s *Maverick*.

**Figure 2.**
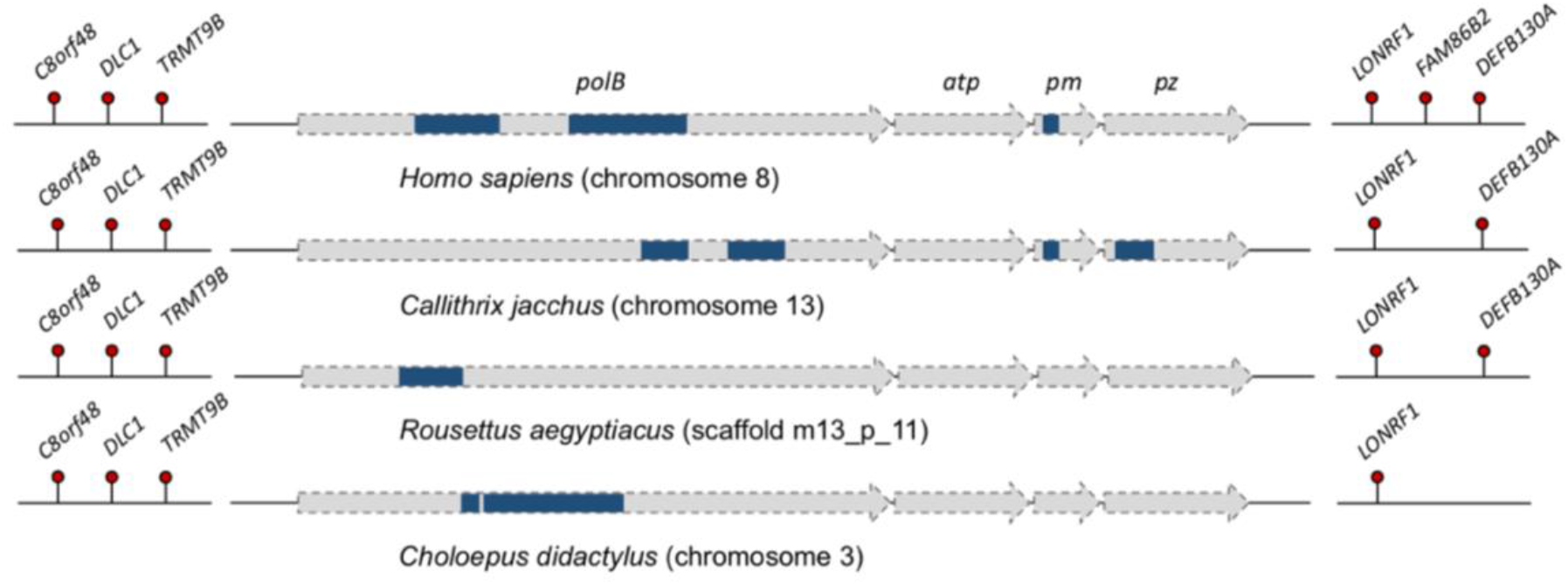
Comparison of the regions orthologous to the chromosome 8 *Maverick* integration in humans (*Homo sapiens*). tBLASTn hits (e-value < 0.05) to the proteins encoded by the Common box-turtle (*Terrapene carolina*) *Maverick* are shown in blue. The arrow outlines represent a segment of the open reading frames of the box-turtle’s *Maverick*.

All the integrations we analysed were highly degraded showing strong signs of being non-functional at the protein level. Firstly, the open-reading frames are interrupted with multiple early stop-codon and frameshift mutations. The elements also lack the terminal inverted repeats characteristic of intact *Maverick* elements. In the case of the human chromosome 8 orthologues, all of the region downstream of the *pz* gene seems to be absent. In addition, the genetic distances of the elements are not significantly different from the distances between non-coding regions in their surrounding genomic neighbourhood (Kolmogorov-Smirnov test, p > 0.99, Figures A1 and A2), which suggests the elements have been evolving neutrally. Interestingly, the chromosome 7/8 *Maverick* orthologues appear to have been lost in some mammals since they could not be detected by either method but were still present in related species. For example, the chromosome 7 integration appears to be absent from the genome of the Ugandan red colobus (*Piliocolobus tephrosceles*) while it is present in other colobine monkeys (*Trachypithecus francoisi*, *Rhinopithecus roxellana*). Similarly, the chromosome 8 integration seems to be absent from the genomes of the grey mouse lemur (*Microcebus murinus*) and Coquerel’s sifaka (*Propithecus coquereli*), despite the *polB* marker being found in the northern greater galago (*Otolemur garnettii*) (this sequence is present in the whole genome alignment and shows a BLAST hit to the PolB of the common-box turtle with an e-value of 5e-4).

The presence of the *polB* marker on both the chromosome 7 and 8 orthologues allowed us to estimate a joint phylogeny with the common ancestor of the elements represented by the root of the tree (Figure 3). The Bayesian estimate for the age of this common ancestor indicates it existed at the end of the Palaeozoic or start of the Mesozoic Eras (Figure 3, Table 1). Although we observed heterogenous rates in the analysis, the inferred rates (mean = 0.0025, IQR = [0.0021, 0.0028]) agree with the overall pan-genomic substitution rates reported previously for mammals: 0.0027 nucleotide substitutions/site/million years (31).

**Table 1.**
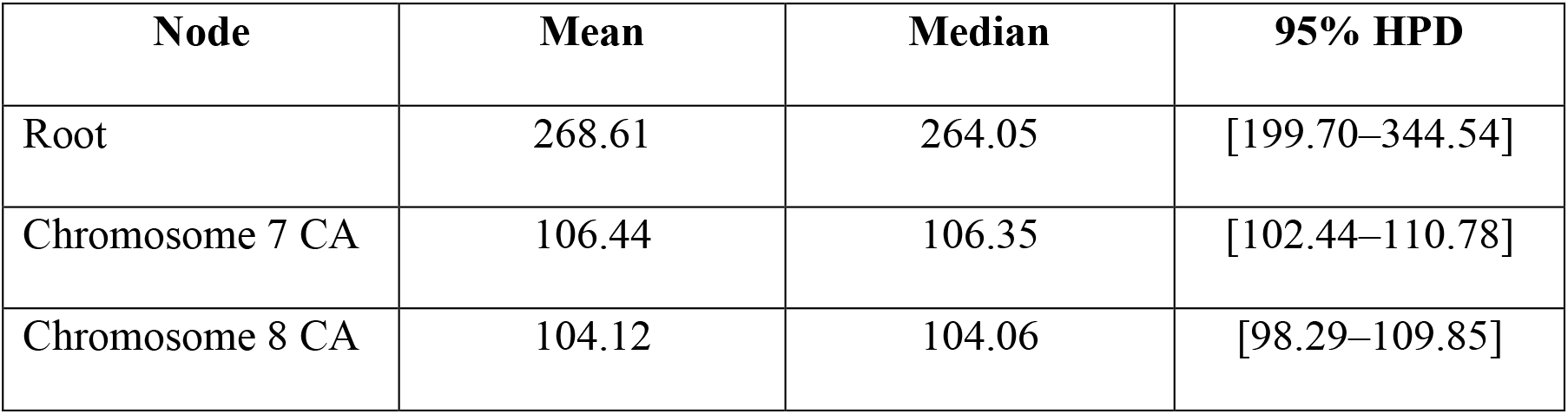
Posterior estimates for the age of the root, and the minimum ages of the chromosome 7/8 integrations (in million years from the present).

**Figure 3.**
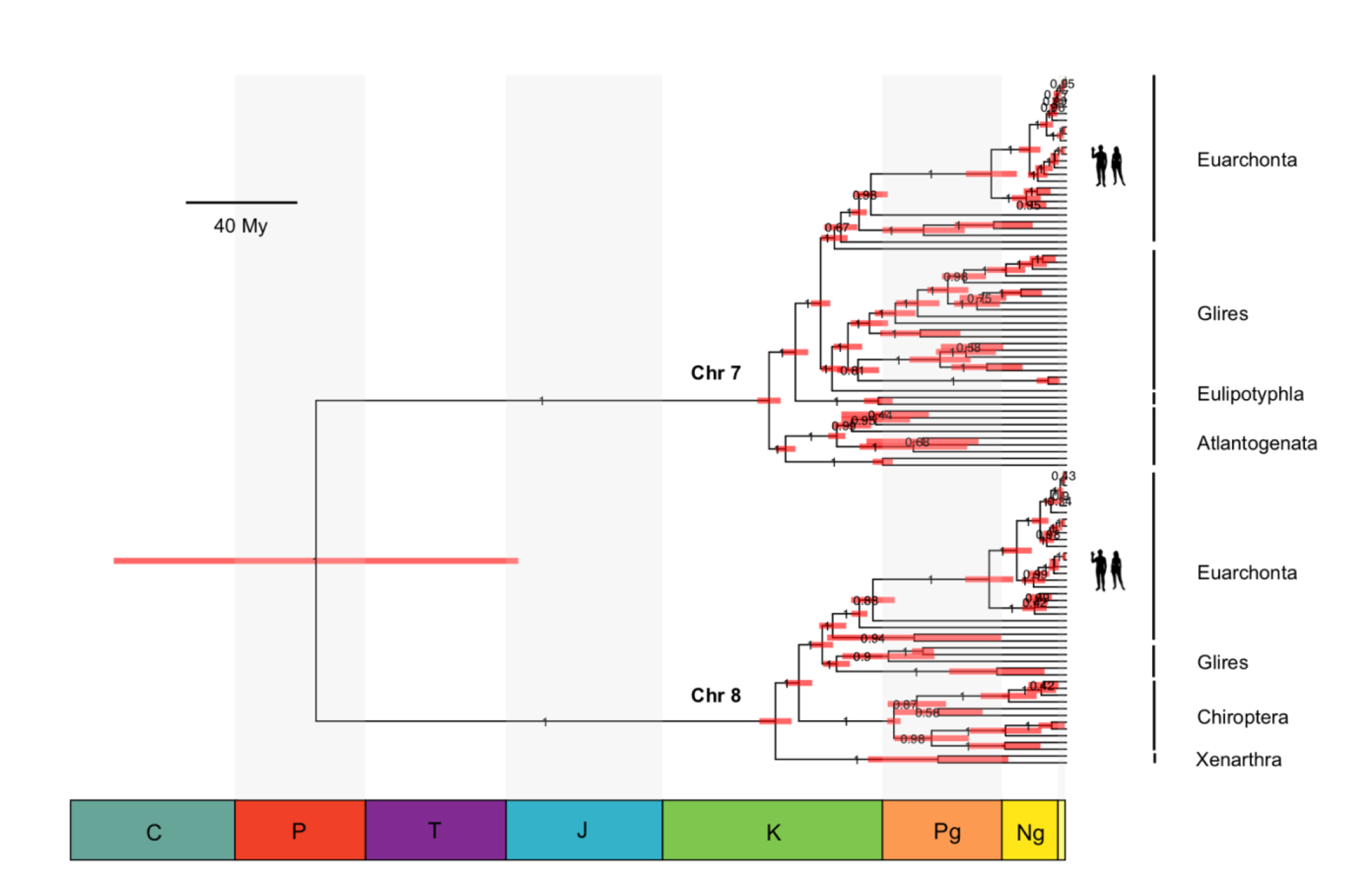
Phylogenetic tree of the mammalian polymerase B-like sequences orthologous to the insertions in human chromosomes 7 and 8. The horizontal node bars represent the 95% highest probability densities (HPD) for the age of the ancestor. Scale bar in million years (My).

## Discussion

Our analyses have revealed that the two *Maverick*-like elements on chromosomes 8 and 7, with an estimated minimum age of 106 (102–111) and 104 (98–110) My, represent the oldest non-retroviral EVEs found in the human genome. An endogenous retrovirus-L element (ERV-L) of a comparable age (at least 104–110 My) has also been identified in humans and shown to be orthologous across the placental mammals (32). Comparison of the sequence divergence between the 5’ and 3’ LTRs of elephant and human (the 5’ and 3’ LTRs would be identical upon integration in the common ancestor), further suggests that the element may have integrated 24-36 My prior to the initial split between afrotherians and boreoeutherians (32). In terms of DNA viruses, an endogenous parvovirus-like element discovered in an intron of the *Ellis van Creveld syndrome 2* gene was shown to be present in primates, carnivores, ungulates and dolphins but not in afrotherians, giving it a minimum age of 98 My (33).

Discovery of the human element found on chromosome 8, together with the one found on chromosome 7, allowed us to gain a more detailed understanding of the evolutionary history of the elements before the diversification of placentals. Indeed, our time-calibrated phylogeny with both elements allowed us to infer the age of the root. This analysis suggests that *Mavericks* circulated in the ancestors of mammals 268 (199–344) Mya, which corresponds to the Carboniferous to Triassic periods. This is supported by the existence of *Maverick polb*-like sequences in the genomes of marsupials and monotremes albeit at non-orthologous positions with respect to the ones we have identified in humans (Table A2). Clear orthology of the chromosome 7 and 8 integrations across placental mammals together with their absence in marsupials and monotremes, suggest that the viruses integrated into the genome of the placental ancestor after the split with marsupials. Molecular estimates place this divergence around 172 (168–178) Mya (34), which is consistent with the age of the earliest fossil eutherian, *Juramaia sinensis*, from the Late Jurassic (160 Mya) of China (35, 36). Thus, it seems likely that active *Mavericks* persisted until the Jurassic Period in the genomes of the early eutherian ancestors.

It also seems that both chromosome 7 and 8 integrations had already been inactivated by the time of the most recent common ancestor of placentals (~105 Mya) (27), pointing to an older age of integration. This is evidenced by deletion of the genes downstream of *pz* in the chromosome 8 orthologues and the absence of the *pz*, *pro* (protease) and *pw* genes in the chromosome 7 insertions. In this respect, these *Maverick* insertions resemble those found in birds which are highly degraded and whose genes contain multiple inactivating mutations (16). We show that the two insertions found in humans and placental mammals are likely to be non-functional since they do not seem to be under selection, they do not localise to piRNA clusters and have been lost altogether on several occasions in several species of mammals.

The reasons for the demise of *Mavericks* in mammals (and birds) remains an open question. Here we hypothesise several plausible scenarios. It has been suggested that *Maverick*s may function as a virophage-induced defence against the infection of large DNA viruses (37, 38), in particular iridoviruses (16). Iridoviruses are important pathogens of fish, amphibians and non-avian reptiles but they do not seem to infect either birds or mammals (39, 40). One possibility is that iridoviruses went extinct in mammals and birds as a result of this *Maverick* defence system. Once their viral hosts went extinct, endogenous virophages (which depend on a host virus for replication) would not have been able to further mobilise and this would have led to their degeneration. This hypothesis could be tested by performing iridovirus infection experiments on cell cultures of hosts which carry intact *Maverick* elements (teleost fish, amphibians or non-avian reptiles). Alternatively, endogenous *Mavericks* could have been co-opted as a defence against exogenous counterparts. Exaptation of endogenous viruses to serve this purpose has been reported extensively, for example in case of the *ev3*, *ev6* and *ev9* genes in chicken (41), *Fv1* and *Fv4* genes in mice (42–45) and the Jaagsiekte endogenous retrovirus (enJS56A1) in sheep which are able to restrict exogenous viruses (46). Evolution of an effective antiviral defence system would lead to the extinction of the exogenous virus and degeneration of the defence locus once the selective pressure from the pathogen has been lifted (47). Therefore, the *Maverick* integrations found in humans and other placentals could represent an antiviral defence system that has served its purpose and is now decaying. However, *Mavericks* have not been linked to any specific pathology so far and their exogenous stage has yet to be observed. Still another possibility is that these *Mavericks* were incidentally inactivated by host innate immune genes that evolved to fight other viral pathogens. For example, cytidine deaminases of the APOBEC3 family function as an antiviral defence in eutherian mammals and have been shown to target both DNA viruses and the DNA stage of retroviruses (48–50).

In addition to the viral arms-race scenarios, other hypotheses relating to changes in host biology could also explain the demise of *Mavericks* in the genomes of birds and mammals. It is possible that the receptor used for entry has been lost/acquired resistance mutations in these animals and the elements could no longer amplify in the germline by reinfection or cross-species transmissions. Support for this idea would require identification of the host cell receptor used for entry of *Mavericks* and assessment of its present state across vertebrates. Finally, the demise of *Mavericks* in these lineages might be linked to terrestrialisation and the origin of the amniotic egg, which may have limited the opportunities for the spread of these viruses and led to their ultimate degeneration. Although the natural history of *Mavericks* is still not understood, it seems they are most successful amongst aquatic organisms which suggests a water-borne mode of transmission (16, 51), perhaps they become active at the early stages when sperm and ova are released into the water by animals with external fertilisation. In fact, potentially active elements reach their highest diversity and copy numbers in fish (where horizontal transfers have also been detected), while only two intact elements have been discovered in amniotes (in turtles and lizards) (16, 17). It therefore seems likely that the reduced opportunities for transmission could have led to a decrease in copy numbers which eventually made the elements prone to extinction.

Our results support a model where *Maverick* endogenous viruses where still circulating in the genomes of the stem mammals at the end of the Palaeozoic and even persisted in the genomes of the eutherian ancestors into the Jurassic Period. The elements found integrated on human chromosomes 8 and 7 represent the relics of ancient viruses that infected our ancestors >102 million years ago. Several hypotheses could explain the demise of *Mavericks* in mammals, which range from exaptation for an antiviral defence system followed by decay, inactivation by an innate immune mechanism, loss/mutation of the host cell receptor or changes to the biology of hosts which lowered their chances of transmission. Experimental studies focusing on the molecular virology of *Mavericks* and the virus-host interactions are needed to shed light on these issues.

## Acknowledgements

This work was supported by the National Academy of Medicine of Venezuela and Pembroke College, Oxford (“Dr. José Gregorio Hernández” Award to J.G.N.B) and a European Research Council Award (101001623-PALVIREVOL to A. K.). We thank Michael Hiller for helping us retrieve the regions of the viral integrations from their whole-genome alignment of 120 mammals.

## Data availability statement

The supplementary materials for this study have been deposited in Figshare (https://dx.doi.org/10.6084/m9.figshare.17819708). The R Code written to compare distributions of pair-wise genetic distances is available on GitHub (https://github.com/josegabrielnb/pair-wise_distributions).

